# Intraspecific Diversity of *Staphylococcus aureus* Populations Isolated from Cystic Fibrosis Respiratory Infections

**DOI:** 10.1101/2024.11.16.623925

**Authors:** Ashley M. Alexander, Hui Qi Loo, Lauren Askew, Vishnu Raghuram, Timothy D. Read, Joanna B. Goldberg

## Abstract

Chronic bacterial infections are often polymicrobial, comprising multiple bacterial species or variants of the same species. Because chronic infections may last for decades, they have the potential to generate high levels of intraspecific variation through within-host diversification over time, and the potential for superinfections to occur through the introduction of multiple pathogen populations to the ongoing infection. Traditional methods for identifying infective agents generally involve isolating one single colony from a given sample, usually after selecting for a specific pathogen or antibiotic resistance profile. Isolating a recognized virulent or difficult to treat pathogen is an important part of informing clinical treatment and correlative research; however, these reductive methods alone, do not provide researchers or healthcare providers with the potentially important perspective on the true pathogen population structure and dynamics over time. To begin to address this limitation, in this study, we compare findings on *Staphylococcus aureus* single colonies versus and pools of colonies taken from fresh sputum samples from three patients with cystic fibrosis to isolates collected from the same sputum samples and processed by the clinical microbiology laboratory. Phenotypic and genotypic analysis of isolated *S. aureus* populations revealed coexisting lineages in two of three sputum samples as well as population structures that were not reflected in the single colony isolates. Altogether, our observations presented here demonstrate that clinically relevant diversity can be missed with standard sampling methods when assessing chronic infections. More broadly, this work outlines the potential impact that comprehensive population-level sampling may have for both research efforts and more effective treatment practices.

**Data Summary:** The authors confirm all supporting data, code and protocols have been provided within the article.

## Introduction

*Staphylococcus aureus* became the most common respiratory pathogen infecting individuals with the genetic disease, cystic fibrosis (CF), in the early 2000s, surpassing *Pseudomonas aeruginosa*. (Cystic Fibrosis Foundation, 2021). Like many CF-associated pathogens, *S. aureus* is a species with significant diversity across lineages (Bernardy et al., 2020; Petit III & Read, 2018; Raghuram et al., 2023) and it has been observed that mixed populations of genetically distinct *S. aureus* strains can co-occur within an individual host (Hofstetter et al., 2024; Langhanki et al., 2018; Long et al., 2020). However, current clinical surveillance techniques generally do not allow for the detection of multiple lineages of *S. aureus* within a single respiratory sample from people with cystic fibrosis (pwCF) as usually only one colony is isolated for sample archiving and testing after selection for specific pathogens or antibiotic resistance (Long et al., 2020). While these methods are useful for identifying key or highly prevalent pathogens that may be virulent or difficult to treat, they are not informative about the true population structure of a given microbe – the number of lineages present and their proportions. In the case of *S. aureus* in particular, clinical processing is heavily focused on identifying whether MRSA (methicillin-resistant *Staphylococcus aureus*) is present in a sample. While this observation is undoubtedly important for informing treatment practices and research alike, it is possible that the identity of a lineage potentially responsible for the virulence or disease outcomes would remain undetected; for example, in the case of infection that contained both MRSA and MSSA (methicillin-sensitive *Staphylococcus aureus*) strains.

We present an in-depth look at the diversity within populations of *S. aureus* isolated from fresh sputum collected from three pwCF by sampling 6-8 single colonies and 1-2 whole-population pool samples. We compare our phenotypic and genotypic analysis of these populations with those of samples processed by clinical microbiology laboratory. For two of three samples, we were able to isolate and identify more than one lineage that made up the *S. aureus* population in the sampled sputum. Altogether, and in the context of the observed phenotypes and patients’ treatment history, our findings show that intraspecific diversity can have significant clinical implications when it comes to traits related to disease severity and treatment potential like antimicrobial resistance, hemolysis, and quorum sensing activity. In the future, methods that account for and quantify intraspecific diversity may assist in the design of more targeted treatment plans for individuals with chronic infections and would inform future research on population structure and dynamics in chronic infections.

## Methods

### Sample collection

The sampling methods used are described in **Figure 1**. Fresh sputum was obtained from 3 different pwCF seen at the Emory Cystic Fibrosis Center. These samples were spread on Mannitol Salt Agar (MSA; Remel) and incubated for 24-48 hours of growth at 37°C. Eight single colonies from each sample were picked and inoculated into 3 mL of lysogeny broth (LB; Teknova). Liquid cultures were incubated overnight at 37°C in a rotating platform before being restruck on *Staphylococcus* Isolation Agar (SIA; BD Difco TSA with 7.5% NaCl) and made into 25% glycerol freezer stocks. All remaining colonies from the MSA pates were scraped together into a ‘pool’ sample which was resuspended in LB media and 25% glycerol and frozen directly. Samples 1_92_SC through 1_97_SC are single colony isolates from patient 1 (CFBR ID 623) and Samples 1_90_Pool and 1_91_Pool were pool samples from the same patient. Samples 2_101_SC through 2_104_SC and 2_106_SC through 2_109_SC are single colony isolates from patient 2 (CFBR ID 196) and its pools from this patient are 2_100_Pool and 2_105_Pool. Samples 3_111_SC through 3_118_SC were single colonies isolated from patient 3 (CFBR ID 311) and 3_110_SC is this patient’s pool sample (**Table 2**). Corresponding clinical isolates, 1_CFBR (patient 1), 2_CFBR (patient 2), and 3_CFBR (patient 3), were collected, processed and archived by the clinical microbiology laboratory and were made available to this study by the Emory Cystic Fibrosis Biospecimen Repository (CFBR) (**Figure 1**).

**Figure 1.**
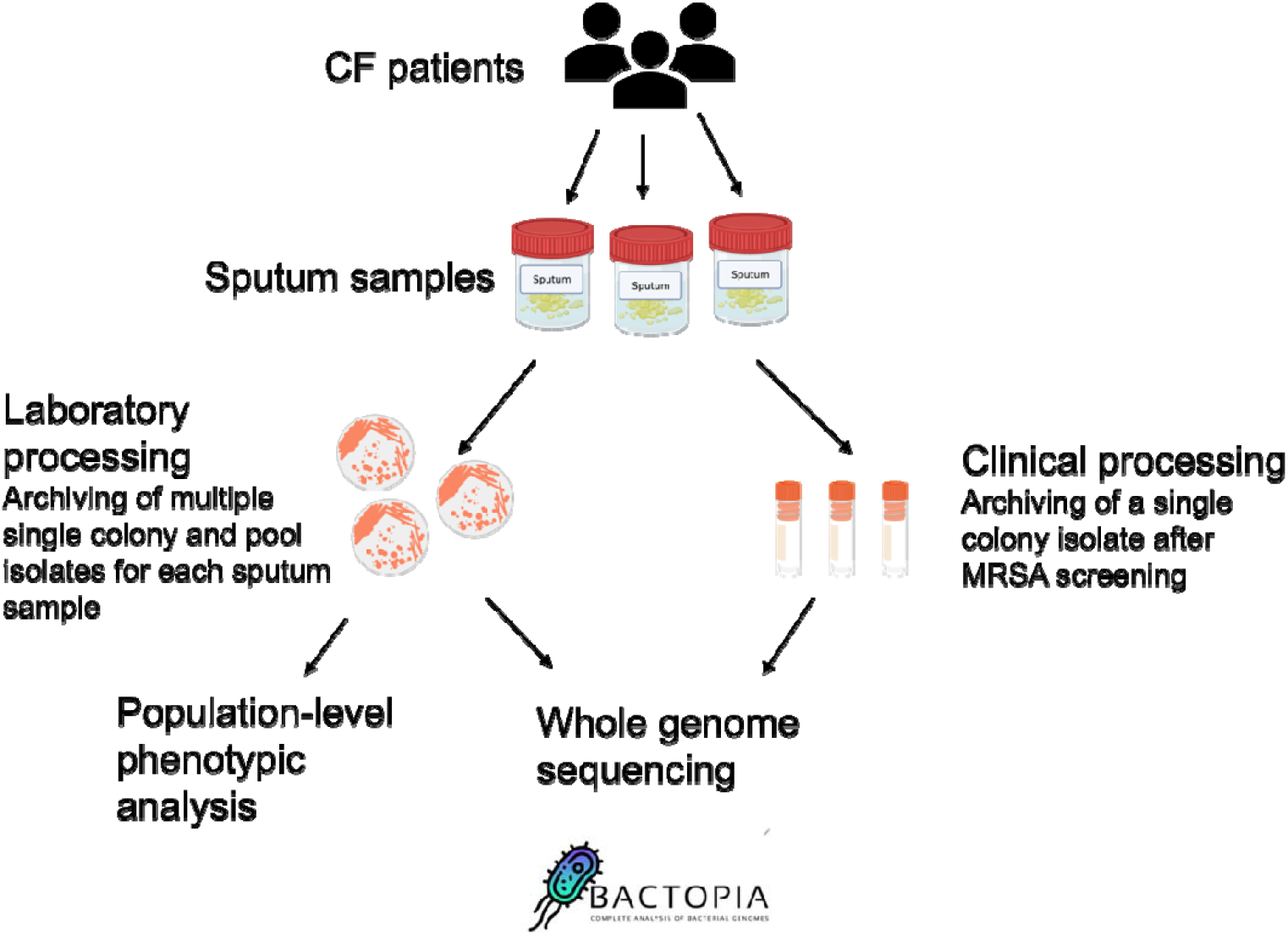
Sampling methods. Method for which isolates were processed from three patients’ fresh sputum samples. 8 single colony isolates were archived per patient sample and 1-2 pool samples were taken for each sputum sample by scraping all remaining colonies. Isolates were then phenotypically and genotypically analyzed for their diversity of traits as described in the text. (Graphic created with BioRender).

### Clinical data collection

Antibiotic resistance testing results of *S. aureus* from isolated from the Emory Clinical Microbiology Laboratory from patients 1 and 2 (CFBR ID 623 and CFBR ID 196, respectively) were made available to us from the Cystic Fibrosis Biospecimen Repository (CFBR) and are outlined in **Table 1**.

**Table 1.** Antibiotic resistance report of *S. aureus* Isolated from the corresponding CF patients from medical microbiology laboratory^a^. Summary of clinical sampling and testing done by the Emory Clinical Microbiology Laboratory from the same sputum samples analyzed in our laboratory.

| Patient | CFBR ID | Sample Date | Hemolysis | Sequence type | Agr group | Antibiotic Susceptibility |  |  |  |  |  |  |
| --- | --- | --- | --- | --- | --- | --- | --- | --- | --- | --- | --- | --- |
|  |  |  |  |  |  | Clindamycin | Daptomycin | Linezolid | Oxacillin | Tetracycline | Trimethoprim/Sulfamethoxazole | Vancomycin |
| 1 | 623 | 2/20/2020 | + | 72 | 1 | Resistant | NA | NA | Susceptible | Susceptible | Susceptible | Susceptible |
| 2 | 196 | 3/12/2020 | - | 225 | 2 | Resistant | Susceptible | Susceptible | Resistant | Susceptible | Susceptible | Susceptible |
| 3 | 311 | 3/12/2020 | - | 5 | 2 | NA | NA | NA | NA | NA | NA | NA |

<sup>a</sup>NA: susceptibility test was not performed by the clinical microbiology laboratory

### Phenotyping

A qualitative analyses of colony morphology was conducted on all isolates including pool samples. While no significant differences in size or texture were observed or recorded, color did vary across samples (white or yellow) and was recorded. Alpha toxin production was observed by noting clear hemolysis on sheep blood agar plates. Congo Red agar (CRA) plates prepared as described by Freeman et al. (1989) were used to determine polysaccharide production levels for all isolates by observing colony texture as previously described (Bernardy et al., 2020; Schwartbeck et al., 2016). MRSA status was determined for all isolates by streaking each isolate and pool sample on ChromAgar™ MRSA plates and observing the presence or absence of purple pigmented colonies, indicating methicillin resistance (Samra et al., 1998).

### Whole genome sequencing

To isolate genomic DNA for sequencing, all isolates and pool samples were streaked on SIA and incubated overnight at 37°C. The next day, cells were collected off plates with an inoculation loop (about a full-loop) and resuspended in 480 µl of EDTA, 20 µl of 10 mg/mL lysozyme and 20 µl of 5 mg/mL lysostaphin were then added to the resuspension and mixtures were incubated at 37°C to lyse cells. After incubation we proceeded with the rest of the protocol outlined for the Promega Wizard genomic DNA purification kit. Sequencing was conducted using the Illumina NextSeq 2000 platform at the Microbial Genome Sequencing Center (Pittsburgh, PA). Genome sequences were evaluated for quality using the program FASTQC (Wingett & Andrews, 2018). Genome sequences were made publicly available at the NCBI BioProject accession <u>PRJNA742745</u>. Samples listed in this BioProject will have the prefix CFBR_EB_Sa before their sample number Sa###, numbers in this BioProject correspond to the same sample number in this study.

### Analysis of Genomic Diversity

Raw fastq files were run through the Bactopia genome analysis pipeline version 3.0.1 (Petit III & Read, 2018). ARIBA outputs were used to determine MLSTs for all samples (Hunt et al., 2017). NCBI AMRFinderPlus version 3.12.8 was used to evaluate antimicrobial resistance associated genotypes (Michael et al., 2019). AgrVATE version 1.0.2 outputs were used to determine the *agr* type for all samples (https://github.com/VishnuRaghuram94/AgrVATE) (Raghuram et al., 2022).

Population-level genomic analysis was conducted by processing fasta files with Parsnp version v2.0.5 to assemble all genome sequences and calculate pairwise single nucleotide polymorphism (SNP) distances using USA300 JE2 NCBI accession NC_007793 as a reference (Kille et al., 2024; Treangen et al., 2014). The program pyANI version 0.2.12 was used to calculate average nucleotide identity (ANI) within patient samples and across all samples (Pritchard et al., 2016). The R-package pheatmap was used to generate a more detailed heatmap of sample ANI values (Kolde, 2012). The program Snippy v4.6.0 (https://github.com/tseemann/snippy) was used to estimate the number of single nucleotide polymorphisms between a previous isolate from patient 2 in 2012 and the isolates obtained in 2020 (Seeman, 2015).

### Experimental evolution of co-isolated strains in the presence of antibiotics

A stepwise serial transfer evolution experiment was designed and modeled after methods outlined in Adamowicz et al. (2020). In order to track the presence of multiple strains throughout the experiment representative isolates 3_112_SC and 3_117_SC were fluorescently labeled by transforming multicopy plasmids pCM29 with GFP (Pang et al., 2010) or pHC48 with dsRed (Ibberson et al., 2016) into either strain via electroporation (Grosser & Richardson, 2016). This resulted the strains were designated 3_112_SC.GFP, 3_112_SC.DsRed and 3_117_SC.GFP and 3_117_SC.DsRed, respectively. For these studies, bacteria were initially prepared by streaking on TSA for single colonies. Individual colonies were then picked and cultured overnight in 3 mL of LB media. Overnight cultures were diluted to an initial OD_600_ culture density of 0.1. Overnight cultures were then diluted to an initial OD_600_ of 0.05 before being inoculated in 96-well plates with antibiotic gradients from 0.25 µg/mL to 128 µg/mL for either oxacillin and vancomycin with two-fold increases across the plate and antibiotic-free control wells. Base-media for all antibiotics tested was CAMHB (Cation-Adjusted Mueller Hinton Broth) or CAMHB + 2% NaCl for oxacillin tests (CLSI, 2018). Mixed cocultures of MRSA isolate 3_112_SC and MSSA isolate 3_117_SC were inoculated at an initial ratio of 1:1 with a final initial OD_600_ of 0.05 for each strain. Plates were incubated for 24 hours at 37 °C with shaking at 450 rpm per growth period.

After each growth period, cells were transferred to a new plate with fresh media and antibiotics. To set up a new 96-well plate during a transfer, 2 µL of antibiotic stock solution was added into each well with 196 µL of media. Culture mixtures were transferred such that 1 µL of culture from wells of each antibiotic concentration was inoculated into a new well of the same antibiotic concentration, and 1 µL of this same culture mixture was added to a new well with 2-fold higher antibiotic concentration, resulting in a stepwise serial transfer method. Transfers were carried out whether or not populations appeared to have growth. This was repeated for 5-6 growth periods. When populations grew in the higher antibiotic concentration, the upper end of the gradient was increased, and the lowest concentration was removed.

OD_600_ and fluorescence readings were taken at the end of each growth period and the 90% minimum inhibitory concentration (MIC_90_) was calculated for each isolate. Growth was defined as an increase in OD_600_ measurement was at least 10% of the antibiotic free control well for a given growth period.

## Results

### Clinical sample analysis

Expectorated sputum was collected from three different pwCF and identical samples were processed, analyzed, and archived by both the Goldberg lab and the Emory Clinical Microbiology Laboratory (**Figure 1**). At collection, the Clinical Microbiology Laboratory performed antibiotic susceptibility testing on single colony isolates from patient 1 and patient 2. Patient 1’s clinically isolated *S. aureus* was identified as being resistant to clindamycin and susceptible to oxacillin, tetracycline, trimethoprim and sulfamethoxazole and vancomycin.

Clinically isolated *S. aureus* from patient 2 was identified as being resistant to clindamycin as well as oxacillin and susceptible to daptomycin, linezolid, tetracycline, trimethoprim and sulfamethoxazole and vancomycin (**Table 1**). Patient 3’s sputum sample was taken shortly before the Clinical Microbiology Laboratory was closed due to the COVID-19 pandemic, therefore they were unable to process this sample for antibiotic susceptibilities. The archived samples were obtained and tested for hemolysis activity; only patient 1’s *S. aureus* clinical isolate was positive for alpha hemolysis. Whole genome sequencing was also conducted on all three samples. Bactopia was used to determine the multi locus sequence types (MLST) for all three samples as well as *agr* group (**Table 1**).

### Phenotypic analysis of populations

To assess the phenotypic diversity of all single colony and pool isolates processed in the Goldberg lab, within and between patients, we first started by assessing the colony morphology of all samples on TSA plates. At initial processing, two small colony variants were selected from Patient 1’s MSA plate, however, upon re-streaking on SIA these isolates were found to be non-viable even after several days of incubation on rich agar. All other isolates from all three patients appeared very similar to each other in colony size and texture. Notably though, colony color or pigment varied across isolates. Patient 1’s isolates were all white in appearance on SIA plates while all of Patient 2’s isolates were yellow. Patient 3’s single colony isolates were yellow while the pool sample 3_110_Pool had both white and yellow colonies, suggesting that there is an additional population (white) that is not represented in any single colony isolate from this patient (**Table 2**).

All isolates from patients 1 and 2 (both pool and single colonies) were positive for alpha toxin hemolysis when cultured on sheep blood agar. In patient 3 only the pool sample and isolates 3_117_SC and 3_118_SC were positive for alpha hemolysis (**Table 2**).

**Table 2.** Compilation of genotypes and phenotypes of single colony and pool Isolates of *S. aureus*^a^ CF Sputum. Complete summary of genotypic and phenotypic testing done on single colony, pool and ChormAgar isolated MRSA isolates from all three patient sputum samples.

| Patient | Isolate ID | Isolate Type | MLST | agr group | MRSA status by ChromAgar<sup>TM</sup> | Colony Color | Alpha toxin Hemolysis | Polysaccharide production on Congo Red |
| --- | --- | --- | --- | --- | --- | --- | --- | --- |
| 1<br>(CFBR ID: 623) | 1_90_Pool | Pool | 72 | 1 | S | White | + | Red/Matte (overproducer) |
|  | 1_91_Pool | Pool | 72 | 1 | S | White | + | Red/Matte (overproducer) |
|  | 1_92_SC | Single colony | 72 | 1 | S | White | + | Red/Matte (overproducer) |
|  | 1_93_SC | Single colony | 72 | 1 | S | White | + | Red/Matte (overproducer) |
|  | 1_94_SC | Single colony | 72 | 1 | S | White | + | Red/Matte (overproducer) |
|  | 1_95_SC | Single colony | 72 | 1 | S | White | + | Red/Matte (overproducer) |
|  | 1_96_SC | Single colony | 72 | 1 | S | White | + | Red/Matte (overproducer) |
|  | 1_97_SC | Single colony | 72 | 1 | S | White | + | Red/Matte (overproducer) |
| 2<br>(CFBR ID: 196) | 2_100_Pool | Pool | 5* 6/7 | 2 | R | Yellow | + | Red/Matte (overproducer) |
|  | 2_101_SC | Single colony | 5* 6/7 | 2 | S | Yellow | + | Red/Matte (overproducer) |
|  | 2_102_SC | Single colony | 5* 6/7 | 2 | S | Yellow | + | Red/Matte (overproducer) |
|  | 2_103_SC | Single colony | 5* 6/7 | 2 | S | Yellow | + | Red/Matte (overproducer) |
|  | 2_104_SC | Single colony | 5* 6/7 | 2 | S | Yellow | + | Red/Matte (overproducer) |
|  | 2_105_Pool | Pool | 5* 6/7 | 2 | R | Yellow | + | Red/Matte (overproducer) |
|  | 2_105_MRSA | ChromAgar Isolation | 225 | 2 | R |  | - |  |
|  | 2_106_SC | Single colony | 5* 6/7 | 2 | S | Yellow | + | Red/Matte (overproducer) |
|  | 2_107_SC | Single colony | 5* 6/7 | 2 | S | Yellow | + | Red/Matte (overproducer) |
|  | 2_108_SC | Single colony | 5* 6/7 | 2 | S | Yellow | + | Red/Matte (overproducer) |
|  | 2_109_SC | Single colony | 5* 6/7 | 2 | S | Yellow | + | Red/Matte (overproducer) |
| 3<br>(CFBR ID: 311) | 3_110_Pool | Pool | 5* 4/7 | 1/2(m) | R | Multicolored | + | Red/Mixed shiny and matte (normal/nonproducer) |
|  | 3_111_SC | Single colony | 5 | 2(f) | R | Yellow | - | Red/Shiny (nonproducer) |
|  | 3_112_SC | Single colony | 5 | 2(f) | R | Yellow | - | Red/Shiny (nonproducer) |
|  | 3_113_SC | Single colony | 5 | 2(f) | R | Yellow | - | Dark/Matte colonies (normal) |
|  | 3_114_SC | Single colony | 5 | 2(f) | R | Yellow | - | Red/Matte and shiny (normal/overproducer) |
|  | 3_115_SC | Single colony | 5 | 2(f) | R | Yellow | - | Red/Shiny (nonproducer) |
|  | 3_116_SC | Single colony | 5 | 2(f) | R | Yellow | - | Red/Shiny (nonproducer) |
|  | 3_117_SC | Single colony | 8 | 1 | S | Yellow | + | Red/Matte (overproducer) |
|  | 3_118_SC | Single colony | 8 | 1 | S | Yellow | + | Red/Matte (overproducer) |

<sup>a</sup>\*, not a complete match; fraction after \* indicates the number of loci that map to that MLST out of the 7 total loci used to assign MLST. +, clear hemolysis; -, no hemolysis. (m) indicates that multiple agr groups were detected. (f) indicates that a frameshift mutation was identified in the agrC locus. MRSA status based on ChromAgar is denoted as resistant (R) and sensitive (S).

All isolates from patients 1 and 2 (both pools and single colonies), were determined to be overproducers of polysaccharide based on their rough and matte appearance on Congo Red Agar (Bernardy et al., 2020). Isolates from patient 3 appeared more heterogenous. Both the patient pool sample and single colony isolate samples presented with some variation on CRA, with isolates 3_110_Pool and 3_114_SC having both textured and shiny red colonies and therefore classified as mixtures of normal and overproducing cells. Isolate 3_113_SC appeared as dark shiny colonies classifying it as a normal producer based on the darker color. Isolates 3_115_SC and 3_116_SC were bright red and shiny on plates classifying them as clear nonproducers. Finally, isolates 3_117_SC and 3_118_SC from patient 3 were determined to be overproducers based on their red and matte appearance (**Table 2**).

ChromAgar™ plates were used as an additional test of MRSA status. Colony growth with purple pigmentation on ChromAgar™ can be reliably used to detect methicillin resistance in *S. aureus* as described in Samra et al. (1998). All isolates from patient 1 showed no growth on ChromAgar™ plates and therefore were determined to be MSSA. All single colony isolates from patient 2 were determined to be MSSA, however both pool samples showed some pigmented colonies on ChromAgar™ plates. The MRSA subpopulation isolated on ChromAgar™ was archived and later included in genomic analyses as sample 2_105_MRSA. Single colony isolates from patient 3 3_111_SC, 3_112_SC, 3_113_SC, 3_114_SC, 3_115_SC and 3_116_SC as well as the pool sample 3_110_Pool were determined to be MRSA with pigmented colony growth observed for all of the above samples. Samples 3_117_SC and 3_118_SC from the same patient were determined to be MSSA based on their lack of growth on ChromAgar™ (**Table 2**).

### Genotypic analysis

Whole genome sequence data was successfully obtained for all isolates. Genomes were assembled and processed using the Bactopia pipeline. Multi locus sequence type (MLST) was determined for each genome using the MLST tool included in the Bactopia pipeline which for *S. aureus* includes variable alleles across 7 loci. To determine isolate relatedness using single nucleotide polymorphism (SNP) distances, assemblies were processed with Parsnp using USA300_JE2 NCBI accession number NC_007793 as a reference strain (Li et al., 2024; Petit & Read, 2020; Treangen et al., 2014). SNP distances computed by Parsnp were then plotted on a phylogenetic tree using ggtree (Yu et al., 2017) (**Figure 2A**). Samples from individual pwCF generally cluster closely together. However, there were exceptions, including 3_117_SC, 3_118_SC and 3_110_Pool which appears to cluster separately from other isolates from patient 3, and 2_105_MRSA and 2_CFBR which cluster separately from other isolates from patient 2. These exceptions reflect other phenotypic and genotypic findings. In the case of patient 3, samples 3_117_SC and 3_118_SC had differences in their MRSA status, *agr* group, sequence type, and hemolysis compared to other single colony isolates and the pool sample 3_110_P. The pool sample from patient 3 likely contained sequence data from both detected populations (MSSA and MRSA) and therefore did not cluster with either group of single colony isolates. For patient 2, all single colony isolates and pool samples clustered together, with the exception of the MRSA isolate 2_105_MRSA. 2_105_MRSA shared some genotypic similarities with the clinical sample 2_CFBR including its methicillin resistance status however, these two samples are not part of a monophyletic group, reflecting their genetic distance. For patients 1 and 3 the corresponding CFBR samples, 1_CFBR and 3_CFBR, respectively, clustered with isolates from the same patients. Samples CFBR_2 and USA_300_JE2 had the lowest ANI value of 98.94% and several samples were identified as 100% identical. Among patient 1 samples, 1_95_SC, 1_91_Pool and 1_90_Pool were all identical with 1_95_SC and 1_91_Pool also being identical to 1_CFBR. From patient 2 samples 2_100_Pool and 2_107_SC, 2_105_Pool and 2_102_Pool, 2_109_SC and 2_102_SC were all identical based on ANI. From patient 3, 3_112_SC was identical to 3_115_SC and 3_113_SC. Overall, ANI analysis confirmed that isolates from patient 1 are genetically identical. Based on ANI, all patient 2 samples clustered closely together including the pool samples again with the exception of the CFBR_2 isolate and the 2_105_MRSA sample. When comparing ANI across isolates from patient 3 samples 3_117_SC and 3_118_SC cluster closely to the reference strain USA300_JE2_Ref but were distant from other patient 3 isolates and distant from other isolates across the sample set and 3_110_Pool did not cluster with either set of single colony isolates from patient 3.

**Figure 2.**
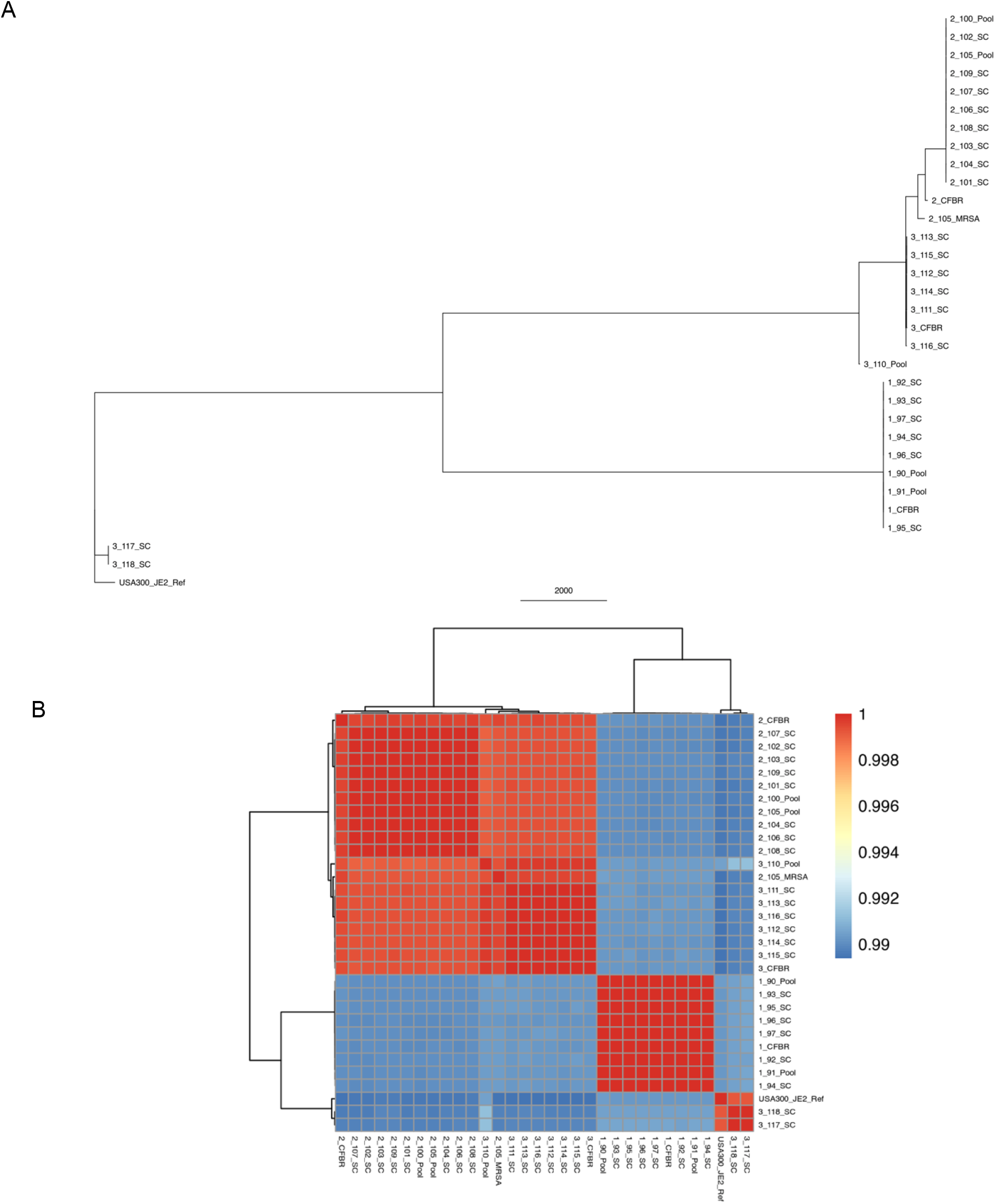
Genetic distances across all patient isolates. A) Phylogenetic tree based on SNP distances computed with the program Parsnp with USA300_JE2 as a reference. SNP distances range from 0 to 14,500. Phylogeny scale (2000) refers to pairwise SNP distance. B) Heatmap generated with the R-package pheatmap visualizing the ANI distances between all patient samples calculated using the python package pyANI. Scale is based on ANI difference (0.00 = 100% identical sequences).

Genes associated with antibiotic resistance were identified in the genomic data for all isolates using the NCBI AMRFinderPlus program as part of the Bactopia genomic analysis pipeline (Michael et al., 2019; Petit & Read, 2020) (**Figure 3**). Patient 1’s isolates all shared the same antibiotic resistance gene profiles. All single colony and pool isolates from patient 2 had identical antibiotic resistance gene profiles while the isolate obtained from the Clinical Microbiology Laboratory and 2_105_MRSA isolates were very similar with the exception of the *sep* allele being present in 2_105_MRSA. All isolates from these patients had alleles associated with resistance to fosfomycin (*fosB*) and tetracycline (*mepA* and *tet38*). While no isolates from patients 1 and 2 had methicillin resistance associated genotypes, *mecA, mec1* or *mecR1* isolates from patients 1 and 2 had variants of *blaI*, and *blaZ* associated with beta-lactam resistance. Additionally, all isolates from patient 1 had alleles *aadD1* and *erm(A)* associated with resistance to aminoglycosides and erythromycin. All isolates from patient 2 and 3 also had the *blaR1* gene associated with beta-lactam resistance - a signature not present in any patient 1 isolates. All alleles associated with antibiotic resistance in the NCBI AMRFinderPlus database with the exception of *sed, sel26, seIX, splE*, and *sep* but including all *mec* alleles associated with methicillin resistance were identified as present in the genomes of all single colony from patient 3 except for 3_117_SC and 3_118_SC. Isolates 3_117_SC and 3_118_SC had genetic signatures associated with fosfomycin, tetracycline and the three *bla* beta-lactam resistance alleles in *blaI, blaR1*, and *blaZ* among antibiotic resistance genes assigned to an antibiotic class. The Pool sample from patient 3 has a unique antibiotic resistance gene profile compared to either group of single colony isolates - missing notably the *mepA* allele associated with tetracycline resistance along with several alleles not assigned to an antibiotic class.

**Figure 3.**
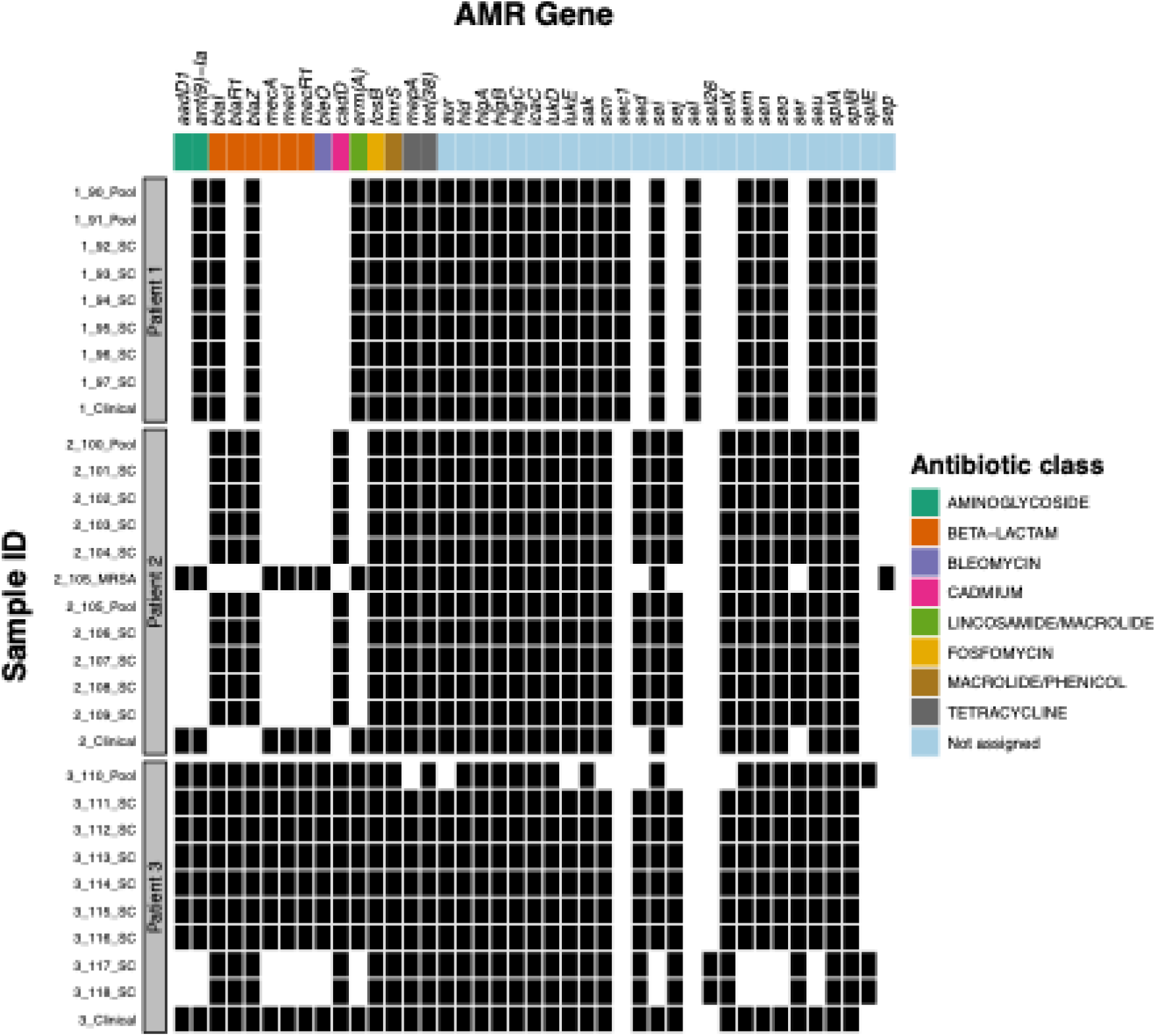
Antimicrobial resistance genetic signatures. Black boxes indicate presence of allele associated with antimicrobial resistance as identified by the AMRFinderPlus tool included in the Bactopia pipeline. Colors across the top of the figure group genes by the antibiotic class their resistance is associated with.

Output from agrVATE (included in the Bactopia pipeline) was used to determine *agr* type and detect any disruptive mutations in *agrC* (Raghuram et al., 2022). All assembled genomes from patient 1 were identified as *agr* group 1 with no frameshifts in the *agrC*. All assembled genomes from patient 2 were identified as group 2 with no frameshifts in the *agrC*. Isolates 3_111_SC through 3_116_SC from patient 3 were classified as *agr* group 2 and having frameshifts present in their *agrC* reading frame; this finding is supported by their lack of hemolysis activity – a trait controlled by the *agrC* (Traber et al., 2008). AgrVATE was able to identify multiple *agr* types present in isolate 3_110_Pool’s coding sequence – the pool sample for patient 3, it was ultimately classified as *agr* group 2. Isolates 3_117_SC and 3_118_SC from patient 3 were identified as *agr* group 1 with no frameshifts in their *agrC* coding sequence (**Table 2**).

### Genomic comparisons with a previously sequenced *S. aureus* isolate

Our group had previously processed a sample taken from patient 2 in February of 2012 (CFBR ID 196; Dr. Eryn Bernardy) which was identified as MLST 225 and MRSA (Bernardy et al., 2020). Using the program snippy we were able to estimate the number of SNPs between the 2012 isolate (Dr. Eryn Bernardy) and an isolate from the 2020 sample set and we found 958 variants (Seeman, 2015). Pairwise SNP distances across recent samples showed that the 2_105_MRSA sample has an average SNP distance of 657 to all single colony isolates from patient 2 but only 294 SNPs compared to the CFBR clinical isolate from the same sampling event 2_CFBR.

### Experimental evolution of co-isolates in the presence of antibiotics

We hypothesized that co-isolated populations may benefit each other’s ability to evolve and persist in the presence of commonly used antibiotics used to treat *S. aureus* infections - oxacillin and vancomycin. Oxacillin is most commonly used to treat MSSA infections while vancomycin is frequently used for treating persistent MRSA infections. To test this, we conducted a stepwise serial transfer evolution experiment where populations were exposed to increasingly higher concentrations of antibiotics over the course of 5 growth periods (**Figure 4A**). We chose representative isolates for the two sampled lineages, 3_117_SC for the MSSA isolate and 3_112_SC for the MRSA isolate. Fluorescence signal was used to identify isolate growth and both color versions (DsRed and GFP) of each isolate was tested in the presence of both antibiotics in monocultures and cocultures.

**Figure 4.**
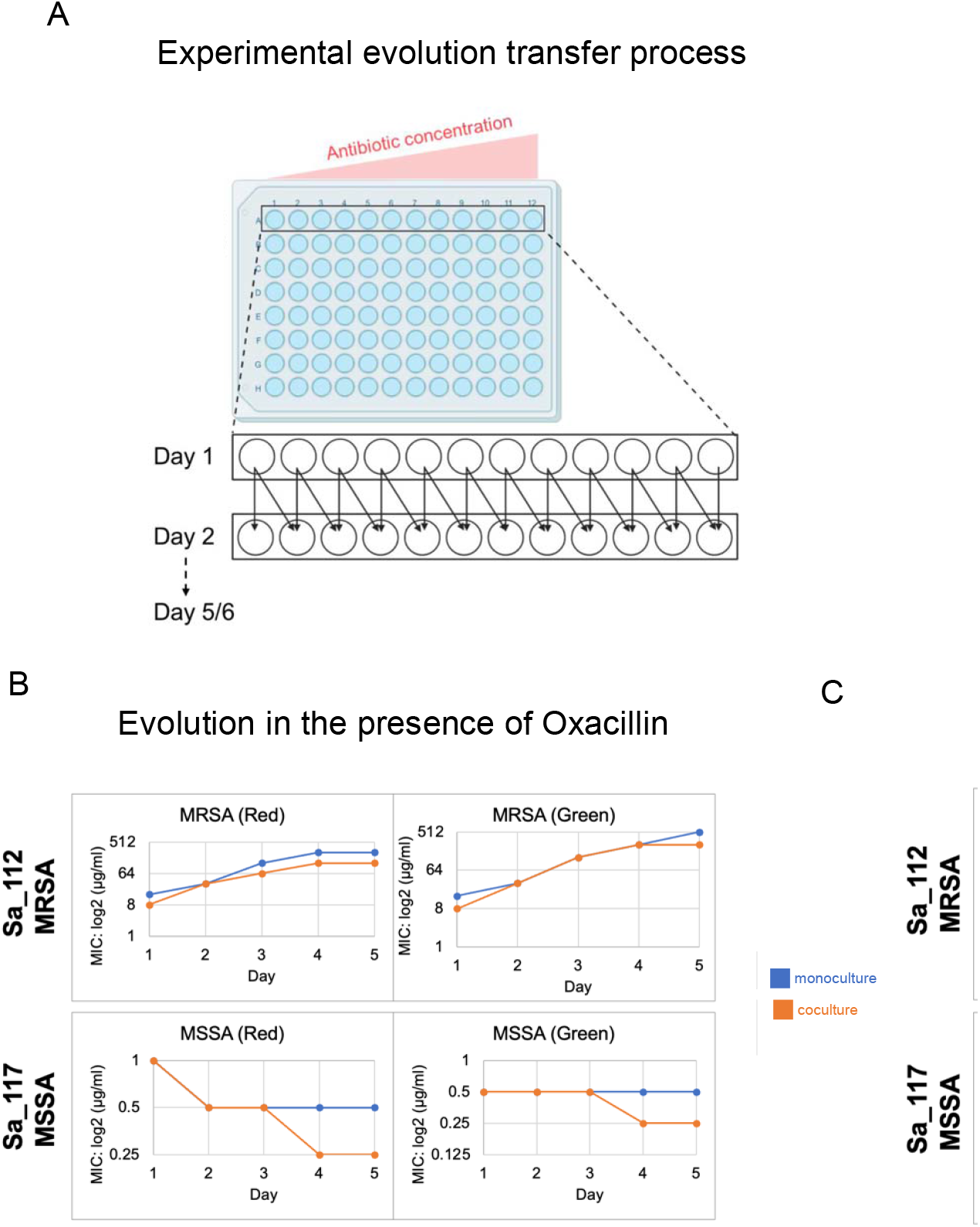
Stepwise experimental evolution of co-isolates alone and cocultured with oxacillin and vancomycin. A) Co-isolates 3_112_SC (MRSA) and 3_117_SC (MSSA) experimentally evolved in monoculture and coculture in the presence of increasing concentrations of oxacillin. Blue lines denote monoculture MIC_90_ values over 6 growth periods and orange lines denote MIC_90_ values of the indicated isolate when in coculture with its co-isolate. Antibiotic gradient increased at 2-fold increments and each fluorescent version of each isolate (DsRed or GFP) was tested.

In the case of oxacillin 3_117_SC (MSSA) did not significantly increase its antibiotic tolerance when evolved alone in monoculture and it was not benefited from evolving in the presence of 3_112_SC (MRSA) as its antibiotic resistance generally did not change and became even slightly more susceptible to oxacillin after evolving in coculture (**Figure 4B**). While 3_117_SC (MSSA) was maintained in the coculture population with no antibiotics present it did appear to be lost from the coculture population at all levels of antibiotics after the third growth period. 3_112_SC (MRSA) initially had an oxacillin MIC_90_ of 16 µg/mL when cultured alone and at the end of the 5 growth periods had reached an MIC_90_ of more than 256 µg/mL. Like 3_117_SC (MSSA), 3_112_SC’s (MRSA) ability to evolve increased antibiotic resistance was not greatly altered by the presence of its MSSA co-isolate 3_117_SC (MSSA) (**Figure 4B**).

When evolved with vancomycin, 3_117_SC (MSSA) began the experiment with an MIC_90_ of 1 µg/mL and achieved an MIC_90_ of at least 4 µg/mL after 6 growth periods. Again, evolving in the presence of its co-isolate 3_112_SC (MRSA) did not greatly alter 3_117_SC’s (MSSA) ability to evolve greater resistance to vancomycin (**Figure 4C**). Despite starting at a lower initial MIC_90_ of 0.5 µg/mL, 3_112_SC (MRSA) was also able to greatly increase its tolerance to vancomycin over the course of 6 growth periods. Unlike 3_117_SC (MSSA) in oxacillin, 3_112_SC (MRSA) persisted in coculture with its more-vancomycin-tolerant co-isolate (**Figure 4C**).

## Discussion

### Sampling single colonies omits the true intraspecific diversity of CF *S. aureus* populations

Our initial hypothesis for this study was that by sampling both single colonies and pool samples of *S. aureus* populations isolated directly from freshly expectorated sputum we could capture intraspecific diversity that may be missed by traditional clinical sampling methods. In support of this hypothesis, we found that the strain composition of two out of three archived clinical samples investigated was different compared to the majority of single colonies or pooled samples that were collected from their respective sputum samples. In the remaining patient sample (Patient 1) where no genetic diversity was detected amongst pooled, single colony samples, or archived clinical samples, morphological diversity was observed in the form of small colony variants however we were not able to further study these variants as they could not be recovered from the archived frozen stocks. Interestingly, we also found that in the case of Patient samples 2 and 3, selection for methicillin-resistant (MRSA) isolates masked *S. aureus* population diversity, as two of our pools contained both MRSA and MSSA strains but only the MRSA isolates were identified by the clinical microbiology laboratory (**Table 1**). Together, these findings suggest that both single isolates and pooled samples should be collected in order to capture the within-species *S. aureus* genetic diversity in a patient.

Specifically, we found that in the case of patient 2, that the population of MSSA (all patient 2 isolates except for 2_105_MRSA) is likely the dominant lineage present in this sputum sample which is not reflected in the archived MRSA clinical sample. In the case of patient 3 an unsampled MSSA lineage was also identified by our sampling methods, which had not been recognized by the clinical microbiology laboratory (**Table 1**). While potentially not as numerically dominant as the MRSA population, this MSSA population showed evidence of intermediate vancomycin resistance based on its high MIC at the start of the antibiotic resistance evolution experiment (**Figure 4**). These MSSA isolates are also positive for hemolysis activity due to their functional *agrC*, while all MRSA isolates from patient 3 have inactive *agrC*, potentially making MSSA the culprit of the majority of *S. aureus* pathogenesis in this patient. Additionally, we noted that patient 3 had recently completed IV treatment with vancomycin at the time of sampling, therefore, the presence of this additional lineage that has intermediate vancomycin resistance in the patient’s sample may have implications for future treatment as well as research aimed at linking patient outcomes and symptoms to pathogen traits. We also suspect that there is a third unsampled lineage present in patient 3’s sample that was not detected by either the clinical lab or the single colony and pool sampling methods. This is based on the white colony morphology which is present in the pool sample 3_100_Pool but is not reflected in any single colony isolate or the clinical sample. The intention of ecological sampling is not to capture the diversity within a system in its entirety, however the method of sampling a single colony along with a pooled sample from an initial selection plate may be a useful approach for both clinicians and researchers similar to what has been suggested in other independent studies (Raghuram et al., 2023).

### Diversity among single colony and pooled isolates yields insight into infection dynamics and pathogenicity

After our initial phenotypic analysis, we had concluded that the *S. aureus* isolates from patients 1 and 2 were nearly identical and that the only within-patient diversity captured in our sample set were the two unique strains identified in patient 3. However, after collecting more data on these samples, a more complex picture unfolded. It was interesting to see that the heterogeneity in patient 3’s samples appeared in the genotypic analysis of 3_110_Pool as well with multiple *agr* types having been identified by the *in-silico* PCR program agrVATE (**Table 2**). ANI analysis also allows us to distinguish the two lineages sampled from patient 3 as truly distinct strains based on the definition outlined in Rodriguez-R et al. (2023), as the ANI values between isolates 3_111_SC through 3_116_SC are all at most 99.00% similar to isolates 3_117_SC or 3_118_SC. Pairwise SNP distances also confirm that the two lineages in patient 3 are unrelated with more than 1400 SNPs between all representative sample combinations. Additionally, we identified an unsampled lineage in our collection existed within patient 2 as both pool samples, 2_100_Pool and 2_105_Pool showed methicillin resistance by ChromAgar™ testing while all single colony isolates were MSSA (**Table 2**). Methicillin resistance was also noted by the CFBR for this sputum sample from patient 2 by the Emory CFBR (**Table 1**). A previous sample from patient 2 from 2012 analyzed by our group had also been identified as MRSA. Based on this previous isolate’s MLST (225) and the number of variants between 2_105_MRSA and a representative isolate from our 2020 sample set 2_102_SC, we concluded that it is not part of the same lineage as the more recent isolates. In fact, the population isolated from the 2_105_Pool ChromAgar™ plate 2_105_MRSA also belongs to MLST (225) and is somewhat closely related to the clinical isolate 2_CFBR taken during the same sampling event with less than 300 SNPs between the two samples. While this is more variation that we would expect to see within a single lineage over 8 years it could be the case that the MLST 225 lineage had already been evolving within this host for some time and was already quite diverse at the time of sampling in 2012. These findings do indicate that there are multiple lineages coexisting within patient 2 and that the MLST 225 population may have been evolving within this patient for potentially decades. Overall, we were surprised at how much diversity we were able to capture with our limited patient sample set and simple isolate collecting methods. The amount of clinically relevant diversity identified in our survey highlights the importance of bringing in ecologically minded sampling practices to patient sample surveys especially in the case of long-term infections like those associated with CF.

### Population dynamics of co-occurring lineages

In the case of the co-isolated MRSA and MSSA lineages, we hypothesized that there may be a benefit for either or both representative isolates 3_112_SC (MRSA) and 3_117_SC (MSSA) when evolving in the presence of oxacillin and vancomycin. This hypothesis was not upheld by our observations and instead it became clear that initial MIC_90_ was a critical factor in an isolates ability to evolve increasing levels of resistance to a given antibiotic. We also observed that growing in coculture with a co-isolate did not appear to have any significant effect on an isolates ability to evolve greater resistance to oxacillin or vancomycin (**Figure 4**).

Taking into consideration the known treatment history of patient 3 (CFBR ID 311), we postulate that our sampling methods have likely captured a shift in the frequency of the cooccurring lineages. It is also likely that it was the MSSA population, which is not reflected in the clinical sample, is more virulent than the MRSA population due to its functional *agrC* and active hemolysis (Traber et al., 2008). Additionally, the dysfunctional *agrC* in the MRSA population could signify its long-term presence in the host as reduced *agr* activity is associated with persistence in chronic infections and immune evasion (Kwiecinski & Horswill, 2020). With this in mind, it is possible that antibiotic treatment combined with elevated host immune activity would keep the more virulent MSSA population in check and the chronic-type MRSA population would continue to persist and increase in frequency. It is interesting to have potentially captured this dynamic period within a single patient’s infection. However, more broadly, our sampling and analysis of these lineages show that an ecological approach to sampling could yield important information about the genetic heterogeneity within a given sample.

Future directions for this work include further testing and application of the sampling approach described here to a larger amount of patient samples, as well as longitudinal patient samples over time. Further investigation of the interactions and dynamics of co-occurring lineages in the presence of antibiotics is also warranted with the opportunity to identify adaptive mutations in evolving populations. Studies analyzing intraspecific diversity within individual patients could illustrate how *S. aureus* populations change over time within a host and in response to clinical treatment. This initial study provides an exploratory but in-depth analysis of diversity across a small sample set. Importantly, we did not seek to replicate the host environment in our analysis of phenotypic or population analysis, and it is very likely that factors from the host environment and immune system would play a major role in changing the outcomes of intraspecific interactions in the host environment. Our findings presented here lay a foundation for a useful workflow and sampling approach that could be applied to a more extensive sample set and aid in answering highly pertinent questions to the clinical treatment of chronic CF infections and within-host microbial ecology and evolution.

## Author contributions

Ashley M. Alexander – Conceptualization, Data curation, Formal Analysis, Investigation,

Methodology, Software, Visualization, Writing – original draft

HQL – Investigation, Methodology, Software, Visualization LA – Investigation

VR – Software, Methodology

TDR – Project administration, Resources, Software, Writing – review and editing

JBG – Conceptualization, Funding acquisition, Project administration, Resources, Supervision, Writing – review and editing

## Acknowledgements

We would like to thank the labs of Dr. Sam Brown and Dr. Stephen Diggle at the Georgia Institute of Technology for sharing with us *S. aureus* populations isolated from fresh sputum. We would also like to thank Dr. Marvin Whiteley for sharing the plasmids pHC48.dsRed and pCM29.GFP. We would like to thank Dr. Katrina Hofstetter for sharing her expertise on the pairwise SNP distance analysis and phylogeny construction.

